# PhenoMIP: High Throughput Phenotyping of Diverse *C. elegans* Populations via Molecular Inversion Probes

**DOI:** 10.1101/857854

**Authors:** Calvin Mok, Gabriella Belmarez, Mark L. Edgley, Donald G. Moerman, Robert H. Waterston

## Abstract

Whether generated within a lab setting or isolated from the wild, variant alleles continue to be an important resource for decoding gene function in model organisms such as *Caenorhabditis elegans*. With advances in massively parallel sequencing, multiple whole-genome sequenced (WGS) strain collections are now available to the research community. The Million Mutation Project (MMP) for instance, analysed 2007 N2-derived, mutagenized strains. Individually, each strain averages ∼400 single nucleotide variants amounting to ∼80 protein coding variants. The effects of these variants, however, remain largely uncharacterized and querying the breadth of these strains for phenotypic changes requires a method amenable to rapid and sensitive high-throughput analysis. Here we present a pooled competitive fitness approach to quantitatively <u>pheno</u>type subpopulations of sequenced collections via <u>m</u>olecular inversion <u>p</u>robes (PhenoMIP). We phenotyped the relative fitness of 217 mutant strains on multiple food sources and classified these into five categories. We also demonstrate on a subset of these strains, that their fitness defects can be genetically mapped. Overall, our results suggest that approximately 80% of MMP mutant strains may have a decreased fitness relative to the lab reference, N2. The costs of generating this form of analysis through WGS methods would be prohibitive while PhenoMIP analysis in this manner is accomplished at less than 1% of projected WGS costs. We propose methods for applying PhenoMIP to a broad range of population selection experiments in a cost-efficient manner that would be useful to the community at large.

## Introduction

The *C. elegans* haploid genome is compact, containing just over 100 Mb, and yet is capable of generating a complex organism with a defined cell lineage (Sulston et al., 1983). Despite our detailed knowledge of this organism, much of its biology remains unclear. At current, only 9,645 Wormbase genes (Wormbase web site, 2019) have phenotype descriptions reported from either variant alleles or RNAi knockdown experiments, suggesting that the function of nearly half of *C. elegans* protein coding genes remain experimentally uncharacterized. Knowledge of where and when a gene is expressed can provide clues to function and many large data sets have elucidated gene expression patterns across embryonic, larval and adult timepoints. Furthermore, multiple techniques have begun to resolve tissue-specific and even cell-specific expression profiles (Boeck et al., 2016; Cao et al., 2017; Gracida and Calarco, 2017; Kaletsky et al., 2018; Warner et al., 2019). However, this information does not directly reveal gene function *per se*.

Forward genetics screens by methods such as chemical mutagenesis, provide a means of recovering alleles that result in a detectable phenotype of interest such as sterility, lethality, or altered reporter expression. These alleles can then be genetically mapped, sequenced, and functionally analysed. In this manner, a specific phenotype can be screened across hundreds of thousands of mutated genomes, thereby querying a very large search space (Brenner, 1974; De Stasio and Dorman, 2001; Kevin et al., 2006). The identification of causal variants across this space can be a laborious process although a variety of methods now exist to aid in the sequencing and mapping of mutant genomes (Doitsidou et al., 2010; Jaramillo-Lambert et al., 2015; Minevich et al., 2012; Mok et al., 2017). In contrast, a reverse genetics screen by RNAi, generates a smaller potential search space by querying a collection of specific gene knock-down targets for a detectable phenotype in a limited number of genetic backgrounds (Fraser et al., 2000; Kamath et al., 2003; Lehner et al., 2006). Consequently, the solution space is relatively well-defined since validated hits require no genetic mapping, although such screens are generally confined to knocking down gene expression rather than necessarily exploring states of altered protein function. Depending upon assay format, an RNAi screen’s throughput can be comparatively less than a mutagenesis screen. Furthermore its effects may be problematic, producing false negatives or weak hits due to incomplete knockdown or false positives from the knockdown of gene families (De-Souza et al., 2019; Fraser, 2000; Parrish et al., 2000). In both screening methods, the ability to score a detectable phenotype can be affected by the presence of redundant paralogs or entire parallel systems that can compensate for a reduced function (for review see (Jorgensen and Mango, 2002)).

Whether because of paralogs or other reasons, phenotypically weak alleles in both screens are potentially missed or simply disregarded. These weak alleles might be mistaken for stochastic variation in a cursory analysis but could provide important insights into function. For example, such weak alleles could produce small changes in developmental timing or fecundity that would affect population fitness (Diaz and Viney, 2014; Perez et al., 2017; Richards et al., 2013; Schnabel et al., 1997). Subtle population-wide shifts in phenotypic fitness require quantitative methods of analysis that go beyond low-resolution phenotype qualifiers such as slow-growth, sterile, or lethal. In recent years, strides have been made in the quantitative analysis of fitness (Crombie et al., 2018; Elvin et al., 2011; Ramani et al., 2012). Advances in next generation sequencing technologies have led to a number of quantitative approaches to population analysis of singular genetic backgrounds by comparing deeply-sequenced samples for changes to transcription, small RNA populations, and heterochromatin (Araya et al., 2014; Boeck et al., 2016; Daugherty et al., 2017; Warf et al., 2012). Leveraging current sequencing paradigms to analyse population fitness would contribute to the process of assigning function to poorly characterized genes or alleles.

To further expand our knowledge of *C. elegans* gene function, we sought to develop an assay that could 1) mimic the allelic diversity of a forward genetics screen but with a smaller solution space much like a reverse genetics screen and 2) generate quantitative data regarding population fitness to assess potential gene function. We exploited the self-fertilizing hermaphroditic nature of *C. elegans* to grow multiple strains in pools without genetic mixing. We also realized that the distinct mutations in each strain could be treated as a barcode to identify and quantify the representation of the strain in the pool. To assay the mutations and thus the representation of each strain in these pools, rather than use whole genome sequencing, which would have been prohibitively expensive, we adapted molecular inversion probes (MIPs) to identify strain-specific variants (Hiatt et al., 2013). We previously used MIPs for the genetic mapping of temperature-sensitive alleles in a collection of *C. elegans* mutant strains (Mok et al., 2017); here we analyse population growth in a multi-generational competitive fitness assay to <u>pheno</u>type by <u>MIP</u>s (PhenoMIP) by quantifying the proportion of each strain in a pool. As a proof of principle, we utilized the Million Mutation Project (MMP) as a source for our strains. The MMP library of 2007 N2-derived mutant strains harbours a variety of coding alleles including potential null alleles across 8150 protein-coding genes, and coding or splice site-altering SNVs across 19,666 genes (Thompson et al., 2013). The phenotypic consequences for many of these variants remain unexplored; we hypothesized that some may play a role in overall fitness. Therefore, we identified unique genetic markers suitable for detection by MIPs for each strain; using these strain-specific MIPs, we effectively generated barcodes for composition analysis of genotypes within a genetically heterogenous population – analogous to methods used in yeast (Hardenbol et al., 2003). We analysed population composition at multiple timepoints, thus determining the relative fitness for each individual strain within a pool, and thereby cataloging the potentially subtle phenotypes of this collection. Our observations suggest that PhenoMIP can identify strains with a range of population fitness phenotypes, including those that may ordinarily be overlooked. Overall, we show that PhenoMIP is a quantitative approach that combines mutagenized genomes that have been previously sequenced and assays them across multiple substrate conditions in a cost-efficient and high-throughput fashion.

## Results

### Molecular inversion probes reliably track multiple strains within a mixed sample

Previously, we demonstrated the usefulness of MIPs as a method to genetically map mutant alleles (Mok et al., 2017). In that study, our empirical analysis of MIP behaviour suggested that their accuracy and precision were highest when identifying smaller subpopulations of variants. Based on this observation, we recognized that the MIP assay could be applied in a large-scale analysis of diverse compositions of strains with complex mixtures of genomic DNA. The mutagenized strains of the MMP collection presented an excellent test set. The MMP strains have, on average, nearly 400 single nucleotide variants (SNVs) per strain, of which, approximately 80 are protein coding changes (Thompson et al., 2013). These strains represent a unique resource for analysing gene function on a large scale.

As a first step we designed a specific set of MIPs to track strain-specific variants (**Figure 1a**). In order to avoid targeting closely spaced variants that might influence the effectiveness of individual MIP assays and because we wanted to preserve the ability to make pools from any combination of MMP and wild isolate strains, we first combined variants from the 2007 mutant and 40 wild isolates strains of the entire MMP project. We eliminated shared alleles, and then chose SNVs separated by a minimum distance of 300 bp. From this list of unique candidate sites, we generated candidate MIP sequences (Mok et al., 2017) and for each strain we identified the highest scoring MIP sequence on each linkage group. From these top six MIPs, we assigned four representative MIPs specific to each strain (**Figure 1b**, and **Supplemental Data SD1**) with the purposes of tracking chromosomal representation in the event of cross-progeny contamination while maintaining minimal reagent costs.

**Figure 1.**
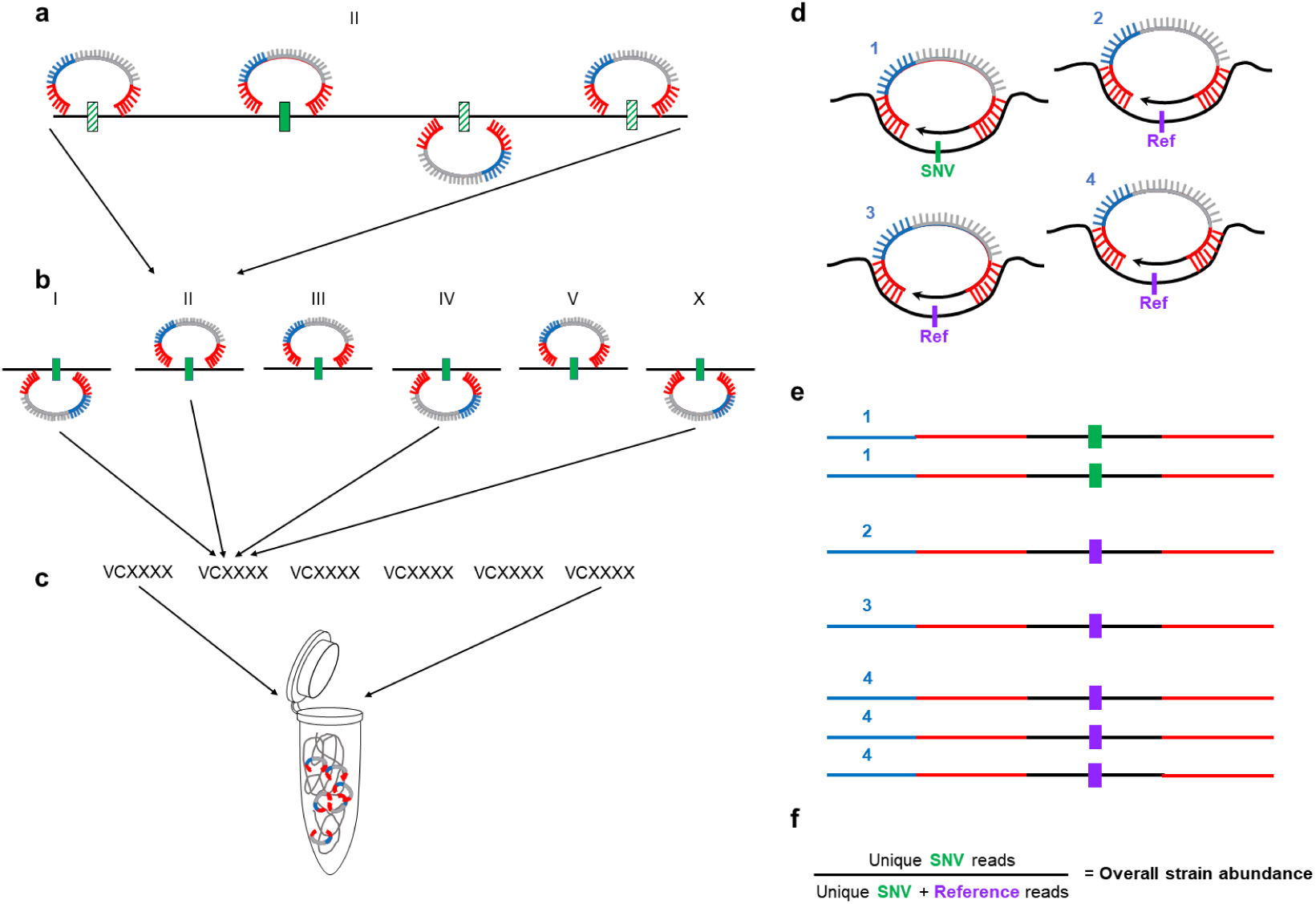
Molecular inversion probes as a system of barcoding *C. elegans* strains. MIP sequences include two annealing arms complementary to target sites (red), a unique molecular identifier (UMI, blue) and a common backbone used for library amplification and barcoding (grey). MIP sites were selected for each of 2047 MMP strains across each chromosome by excluding shared variants from all strains and then choosing sites (regardless of strain) across the genome that were separated by a minimum of 300-350bp. (a) For each strain, MIP candidate sequences were scored (solid and hatched variants). (b) The highest-scoring MIP on each chromosome (solid green) was identified. (c) Four of the six MIPs were then selected to identify a target strain amongst a pool of strain-specific MIPs. The MIPs would therefore have two identifiable states from the gap-fill segment of a sequencing read (d); either the strain-specific single nucleotide variant (SNV, green), or a sequence identical to the reference genome (purple). (e) After sequencing, each sample was demultiplexed by MIP target and further by the UMI to count the total number of unique annealing events specific to the SNV or reference sequences. (f) Values were compared to estimate the percentage of SNV events versus the total annealing events.

To ascertain the representation of each strain in a pool, the four MIPs representing each target strain within a desired composition of strains were combined into a single pool (**Figure 1c**) and used in the generation of MIP sequencing libraries. The libraries were sequenced, demultiplexed and individual annealing events tracked by the unique molecular identifier (UMI) present on each oligo (**Figure 1d, e**). Probe sets were then combined to determine mean relative abundance for each target strain within a pooled set of genomes (**Figure 1f**). To successfully analyse mixed populations in an efficient high-throughput manner the PhenoMIP approach would require 1) a relatively balanced distribution of reads for each probe; 2) a low false positive rate to determine a reasonable lower bound on probe accuracy; and 3) precision between strain-specific targets to ensure that subpopulation analysis was consistent.

To test the above parameters, we generated a pool of 192 MIPs designed to target SNV sites for 48 MMP strains (**Supplemental Data SD2**). We generated five different sets of genomic DNA mixtures composed of subsets of 46 of the 48 MMP target strains in different proportions (two strains failed to yield adequate amounts of DNA) and used these as template samples for the generation of MIP sequencing libraries (**Supplemental Data SD2**). From these libraries we observed the expected composition and proportion of genotypes for the original genomic templates, suggesting that overall cross-MIP interference from multiplexing was negligible (**Supplemental Figure S1a**) and that the variant information from sequencing was correct. We analysed the total number of UMIs for each MIP to gauge the efficiency of each probe. We observed eleven MIP targets that, across all libraries, consistently produced UMI counts below 20% of the mean number of UMIs per MIP in an individual library; these were removed from further analyses (**Supplemental Figure S1b**). To investigate the read distribution of this adjusted dataset, we normalized the UMI counts for each MIP against the minimum read number within its sequencing set. The normalized distribution of reads spanned across a ∼9-fold range with an inter-quartile range of 2-fold to 6-fold suggesting that our distribution was relatively unimodal and ranged within a single order of magnitude (**Supplemental Figure S2a and S2b**).

Next, for each sequenced library, we analysed the MIP reads from target strains that were excluded from the genomic template, calculating a total false positive rate of 1.6×10^−4^ across five MiSeq-generated data sets for which the mean UMI count per MIP was 1630 with 1.2×10^6^ unique capture events across the total set. We also compared two sequencing runs of the same PhenoMIP library with false positive rates of 1.49×10^−4^ at 3.9×10^5^ total capture events versus 1.18×10^−4^ at 5.14×10^6^ total capture events. Combining all data sets we confirmed a total false positive rate of 1.25×10^−4^ across all MIPs. We estimated the mean false positive rate per individual MIP to be 1.29×10^−4^ ± 1.38×10^−4^, which compares well with our prior observations (Mok et al., 2017).

When initially planning experimental design, we chose to work with pools of approximately 50 strains per set, resulting in an expected average initial population abundance of 2×10^−2^. With such a low starting abundance it was important to assess the precision between each set of strain-specific MIPs to ensure that the variation between these probes was low enough to consider their mean value a consistent assessment of strain abundance. We observed the mean standard deviation across all strain-specific MIP sets was 2.33×10^−3^ ± 6.88×10^−3^. Confirming prior observations, the absolute variance between strain-specific MIPs was dependent upon relative abundance within the sample. Subsetting the data, target strains above 5×10^−2^ abundance had a combined standard deviation between strain-specific MIPs of 1.62×10^−2^. Samples with abundance below 2×10^−2^, however, had a combined standard deviation between MIPs of 2.12×10^−4^, which is similar in magnitude to our false positive rate. These findings were in line with our expectations from prior modeling of MIP behaviour (Mok et al., 2017) (**Supplemental Figure S2c**).

From our analyses, we concluded that relatively consistent and balanced pools of MIPs could be generated for future analysis on complex populations; that our false positive rates remained in line with previous observations; and that overall variance among MIPs for a specific target strain was low, especially in the lower ranges of abundance. In combination with our MIP-MAP data (Mok et al., 2017), our analysis conservatively suggests that MIPs can accurately detect variant abundances as low as five standard deviations above the estimated false positive rate. We determined that relative abundances as low as 8.2×10^−4^ would have a high probability of being true signal as our largest false-positive value from the dataset was 7.4×10^−4^. For simplicity, we designated 1×10^−3^ as the minimum abundance required to be considered as biologically present within a given pooled population. Practically speaking, based on an average pooled experiment of 50 strains, this translates to detecting a 20-fold decrease from the expected initial abundance for a target strain. The cut-off value of 1×10^−3^ was the foundation for later analysis of our data sets with these and other MIP pools (**Methods**).

### MIPs identify strain fitness defects over multiple generations

Confident of the estimation capabilities of the MIPs, we selected sets of MMP strains to pool for growth analysis. Each pool was made up of 45-60 different MMP strains and 8-10 independent replicates were grown for multiple generations to look for differences in fitness between the strains (**Table 1**). In addition, to investigate the effects of different propagation methods, three food sources (*E. coli* strains HT115, NA22 or OP50) were used in different experiments and in one experiment two different methods of transfer were used (see below). The proportion of each strain in the pool was assayed at the start, terminal and various intermediate points. To ensure that a similar number of animals was present at the start and in each of the replicates (and different conditions in experiments where more than one condition was assayed), we hand-picked 20 animals from each strain at either the L1 (pool M1, M3, M5) or L4 (pool M7, M8, M10, M11) stages to duplicate *E. coli* seeded plates. We grew these “starter” pools to starvation and combined uncontaminated plates for an estimated 300-700K animals. This population was collected and aliquots containing 5-10K animals were used to inoculate replicate cultures under their specific conditions. Cultures were grown to starvation (72-96 hours at 20-22°C; about a generation) and aliquots transferred to fresh plates. For all pools except M11, animals were transferred by chunking, while M11 replicates were split into two groups with transfer either by chunking or by washing (**Table 1, Methods**). This inoculation-to-starvation cycle was repeated 4-9 times, depending on the experiment. At each cycle a fraction of the population was saved for later DNA analysis. (**Figure 2**).

**Table 1.**
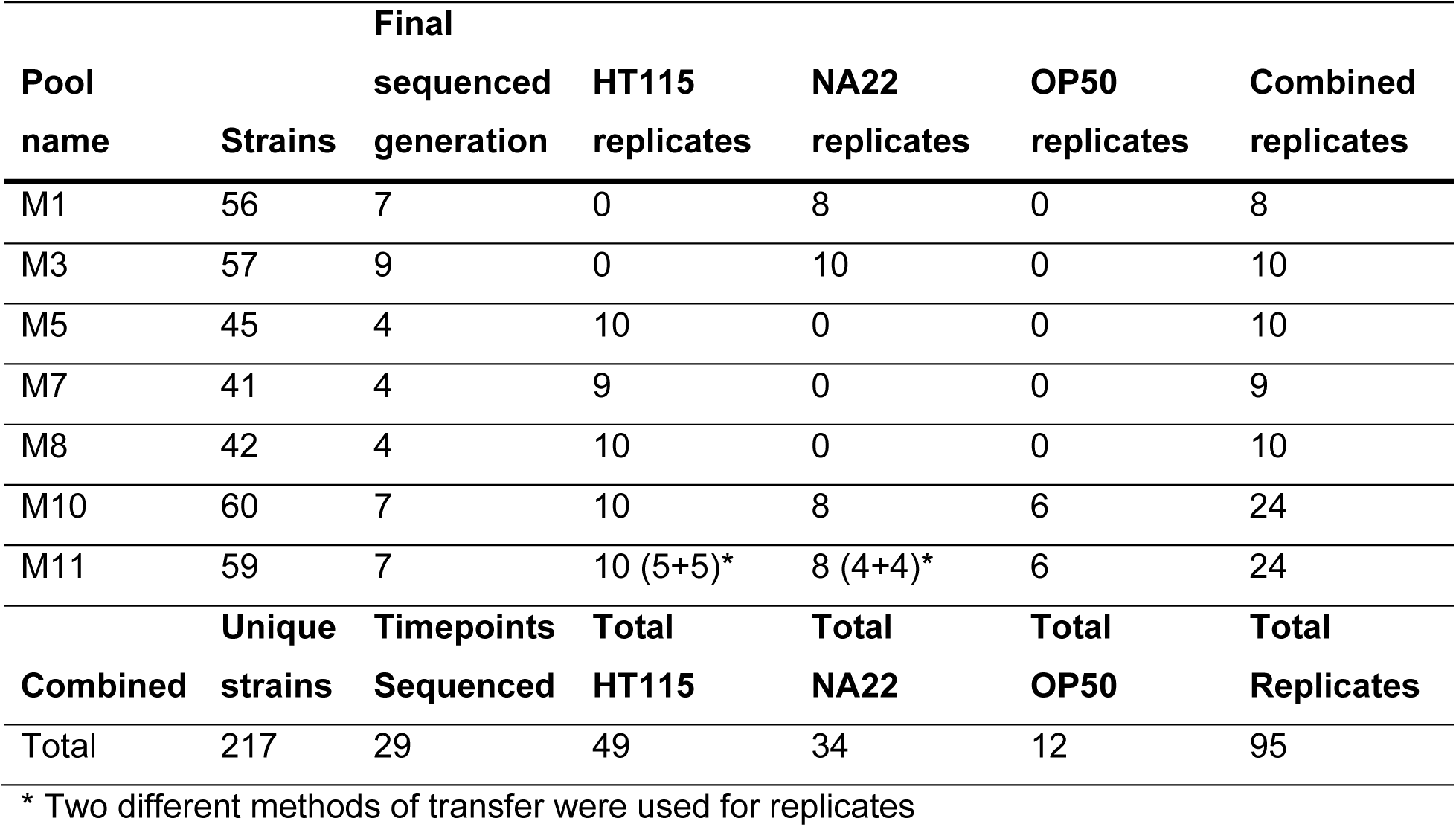
Summary of pooled strains.

**Figure 2.**
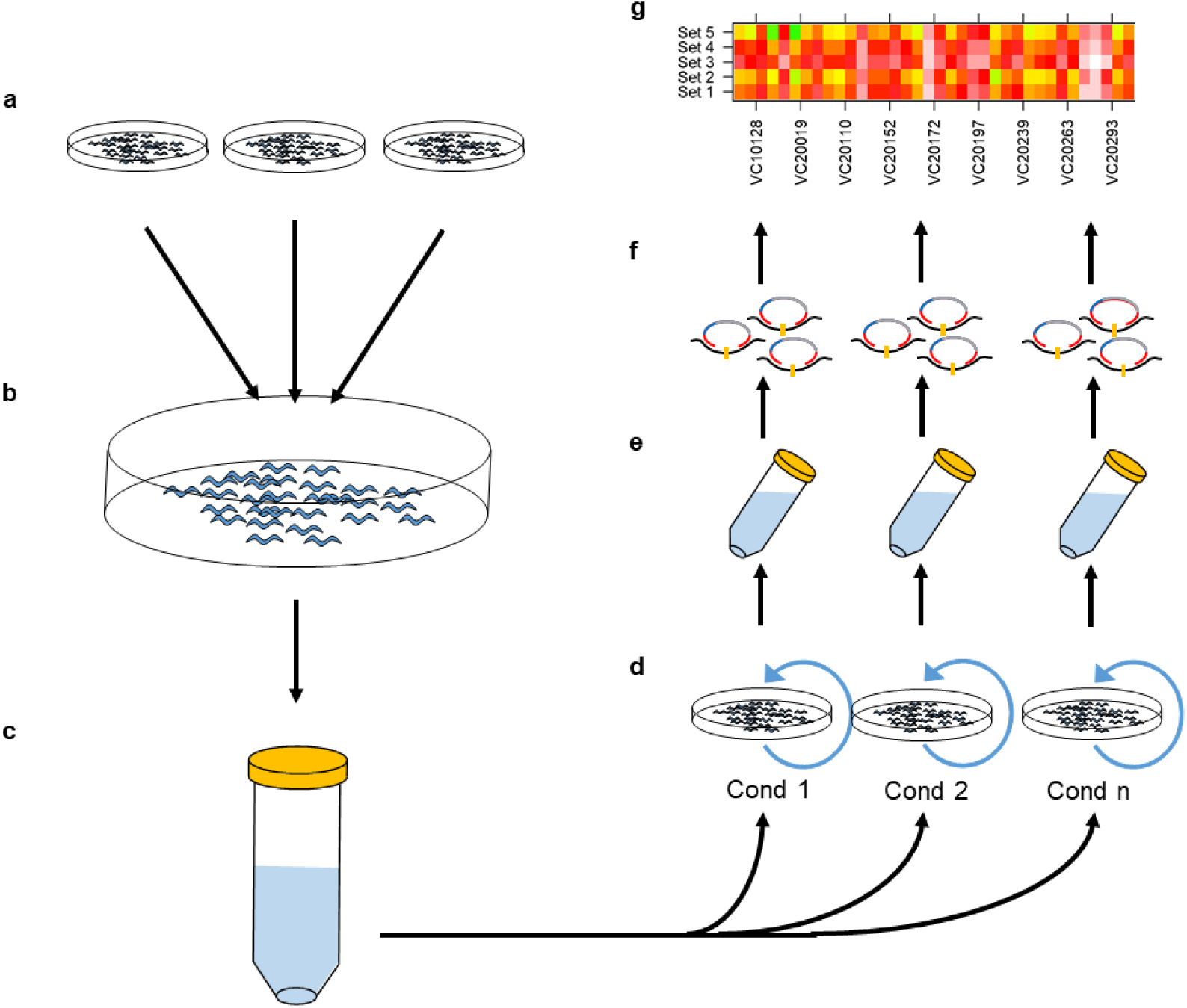
Workflow of PhenoMIP multigeneration competitive fitness assay. (a) MMP strains were selected and grown as separate populations in relative synchronization before 20 animals of each strain at the L1 (pools M1, M3, M5) or L4 stage (M7, M8, M10, M11) are transferred (b) to a communal NGM plate seeded with a bacterial lawn. The communal plates are grown in duplicate until the population has starved. (c) Uncontaminated plates are then washed and combined into a single starting population and counted for population density before being redistributed (d) onto multiple 150 mm NGM plates of varying conditions. Every 72-96 hours, the plates reach starvation and a subpopulation of animals is transferred to a new plate of the same experimental condition. (e) The remaining animals are collected for extraction of genomic DNA to generate MIP libraries for sequencing (f) and data analysis (g) of strain abundance and relative fitness.

*In toto*, we used 217 MMP strains across seven experimental pools (**Table 1, Supplemental Data SD3**) to assay their relative fitness. To check the reproducibility of the data and observe overall trends we applied principal component analysis to the datasets. For example, with the M11 dataset, replicate samples with the same food source and transfer method tended to cluster tightly, but with clusters from different generations separating well after the first generation, particularly along the axis of the first principal component (**Figure 3a, b and Supplemental Figure S3**). Samples also separated by the methods of transfer. PCA analysis on all the M11 samples at a single timepoint shows the effect of food source as well as method of transfer (**Figure 3c and Supplemental Figure S4**). The OP50 replicates were not as well-correlated, and it was observed that these populations starved more quickly than other food sources. Our observations suggest that under a given experimental condition, population composition was changing with each generation in a consistent manner that was detectable by PhenoMIP analysis.

**Figure 3.**
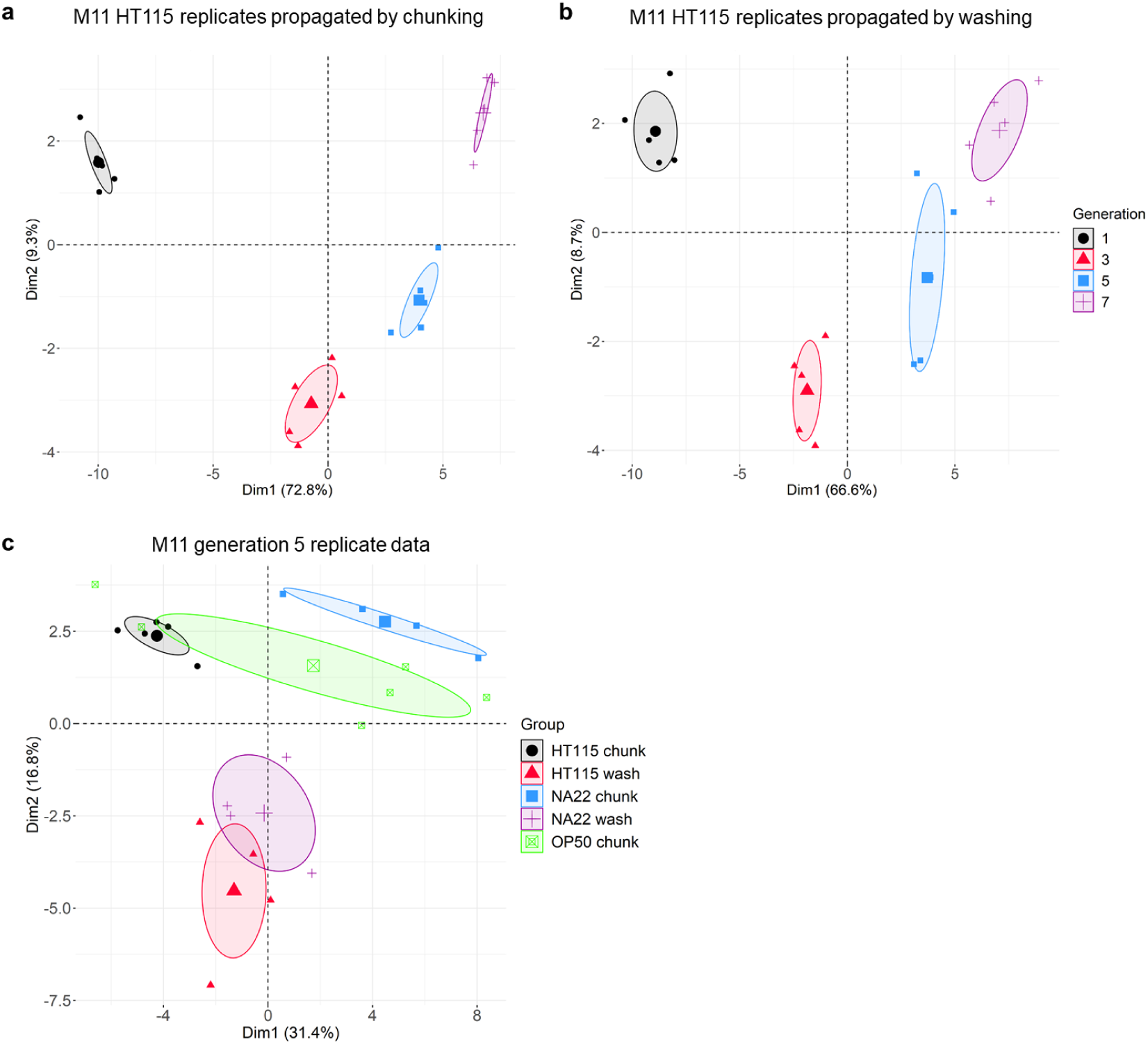
Principal component analysis of PhenoMIP data suggests consistent population stratification related to growth conditions. (a) PCA of M11 HT115 population replicates propagated by chunking and (b) M11 HT115 population replicates propagated by washing are projected along principal component 1 and 2 with samples coloured by generation. PCA of all M11 replicates from generation 5 projected along principal component 1 and 2 with samples coloured by combined food source and transfer method.

Confident that the assay was behaving well overall, we next assessed each strain separately for relative changes in its abundance over multiple generations across multiple replicates. For each replicate condition within a pooling experiment, this effectively created a growth profile for each strain consisting of the total fold-change and the mean fold-change rate (FCR) per generation. For example, **Figure 4a** plots the relative abundance of strain VC20019 in the M11 pools under various conditions. The log-fold change is modest, with the mean across all conditions almost zero, indicating that this strain is of average fitness. Closer inspection suggests that some of the variation is due to the different growth conditions used in M11, with replicates grown on NA22 and transferred by washing showing better than average growth, whereas growth on HT115 and chunk transfer grew less well. In agreement with the overall PCA analysis, growth on OP50 resulted in the most variable log-fold change. We combined results across replicates for all strains to analyse FCR as a distribution across conditions (**Figure 4b and Supplemental Figure S5**). We identified 15 strains that failed to thrive (class 0) in the initial pool expansion steps (initial abundance < 2.5×10^−3^) suggesting they harboured potentially strong deficits to population fitness (**Supplemental Table S1**). We classified the remaining 202 strains using 393 sequencing libraries across seven competitive fitness pooling experiments on 95 replicate conditions to generate profiles for 170 strains grown on the bacteria HT115, 149 strains grown on NA22, and 105 strains grown on OP50 (**Supplemental Figure S6a**). While we observed more subtle differences within some strains for growth on different bacteria and even for methods of transfer (**Supplemental Figure S6b,c**), we observed pronounced differences in growth profiles between strains and focused further analysis on this feature. We observed strains that exhibited poor growth with steep population decline suggesting fitness defects as well as strains with enhanced growth when compared to our reference strain VC20019. Based on these observations, we classified each strain into one of four classes as determined by its mean FCR across all experimental replicates (**Table 2, Supplemental Data SD3**). Classes were designated using a simple 10-generation growth model to calculate a final abundance (A_10_) based on the log_2_-transformed mean fold-change rate 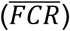 such that

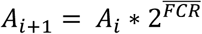

**Table 2.**
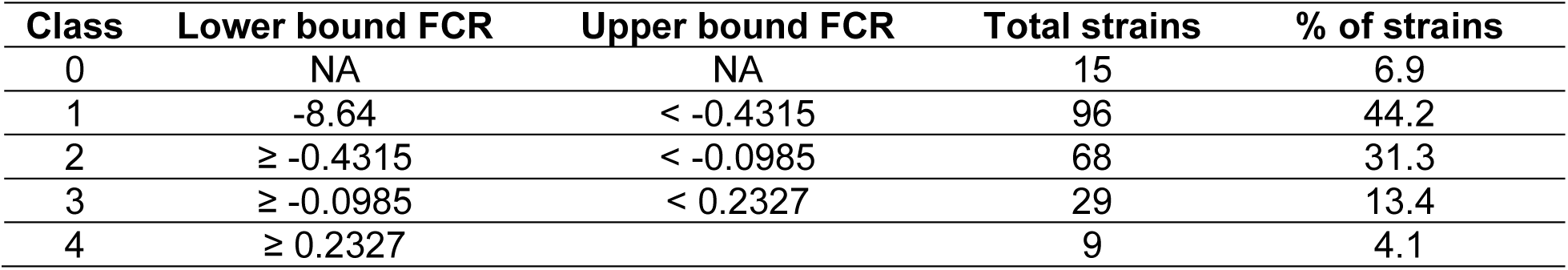
Mean fold-change rate summary.

**Figure 4.**
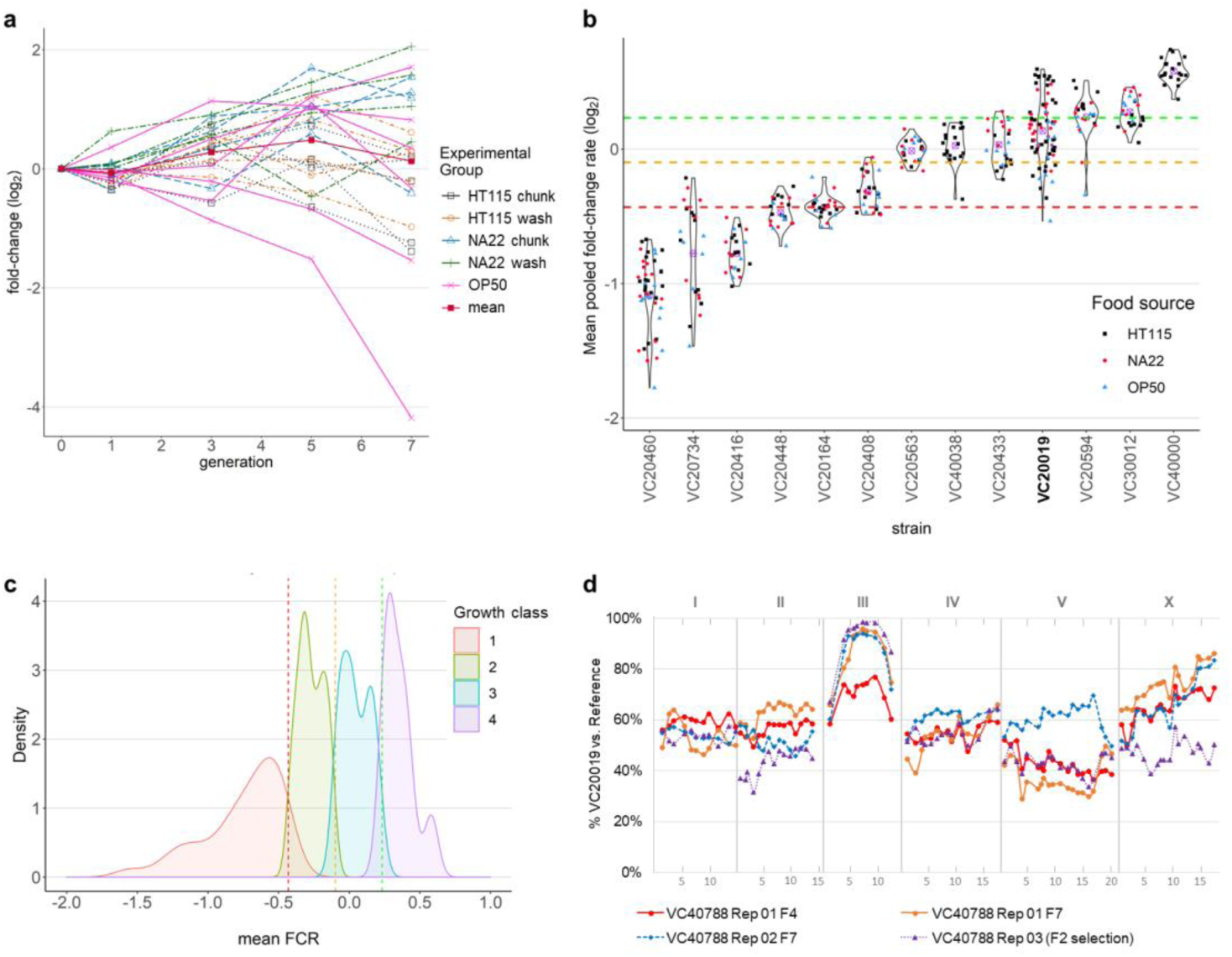
Relative fitness can be quantified by PhenoMIP and classified into subgroups. (a) Line graph of VC20019 growth rate from pool M11 with y-axis showing fold-change (log_2_) in abundance relative to initial abundance at generation 0 (starting population) across multiple generations (x-axis). Replicates are coloured by experimental food source and transfer method: HT115 chunk (black squares), HT115 wash (orange circles), NA22 chunk (blue triangles), NA22 wash (green cross), OP50 chunk (pink x) and mean (mean fold change abundance across all replicates, red square). (b) Violin plots of mean fold-change per generation for a representative panel of strains. Each point represents the mean fold-change rate calculated from multiple timepoints for an experimental replicate across one or more pooling experiments. Dots are colour-coded by experimental condition for growth on either HT115 (black squares), NA22 (red circles), OP50 (blue triangles) *E. coli* as a food source with overall mean fold change rate (FCR, purple cross). Coloured dotted lines represent category boundaries using an FCR of -0.4315 (red), -0.0.985 (yellow), and 0.2327 (green). VC20019 (bold) is provided as a reference for comparison to growth rates shown in (a). (c) 202 strains were assigned a mean FCR and subdivided into one of four growth classes with kernel density plots for each class. (d) Mapping data for VC40788, a strain observed to have poor growth rate, identified an interval of interest at III:7.6-10.8 Mb. Mapping was accomplished using two replicates by competitive fitness for wild type growth (orange circle and blue diamond) as well as by identifying F2 homozygous wild-type F2 recombinants in a bulk segregant assay (purple triangle). X-axis units are in megabases across each chromosome.

From our initial modeling of MIP behaviour, we determined a lower limit of 1×10^−3^ on abundance within a pooled sample; we, therefore, used A_10_ cut-offs of 1×10^−3^, 1×10^−2^, 1×10^−1^ as boundaries for determining classes 1 through 4 (**Supplemental Figure S5**). In particular, we observed that the MMP strain VC20019, which we had previously observed as having a rate of growth similar to the lab reference strain N2, fell into class 3 with a 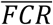 of 0.135 or growth multiplier 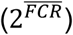 of 1.10 per generation (**Figure 4b**). Subdividing VC20019 data by experimental pool, however, suggested there was potential for pool-specific variation on a larger scale (**Supplement Figure S7**). The higher 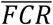 for pool M8 is likely a result of over-representation in the seeding population by double as VC20019 was also conspicuously absent from the M7 seeding population, which was pooled in parallel to M8. Our analysis of the 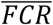 across all strains suggests a wide range of fitness phenotypes across the MMP collection (**Figure 4c**).

### The reduced fitness phenotypes of MMP strains were mapped to candidate mutations

Based on the results of our growth analysis, we hypothesized that underlying mutations within some strains could account for the observed growth rates. We proceeded to genetically map a subset of class 0 and class 1 strains as they exhibited the greatest reduced fitness in comparison to our control strain VC20019. We used our MIP-MAP protocol (Mok et al., 2017) to competitively select against the reduced fitness phenotype and identify a small genomic region containing the associated causal variant. Briefly, mutant strains were crossed with males of the mapping strain VC20019 and the population was grown until starvation. A small portion of the population was then transferred to OP50-seeded 10cm NGM plates. This transfer was completed approximately once per generation for up to 6 generations. Samples were taken at each transfer step and used to prepare genomic DNA for MIP-MAP libraries and sequencing.

We chose five class 0 and two class 1 strains to map, and successfully identified a single locus linked to a reduced population fitness for six strains (**Table 3 and Supplemental Figure S8**); a seventh strain appeared to have two loci. After phenotyping individual strains for possible causes of fitness defects, we were able to assign candidate alleles based on genes with shared phenotypes. In particular, we verified the mapping results of strain VC40788 by following a partially penetrant maternal-effect embryonic lethal phenotype (**Figure 4d**). From VC40788 and VC20019 cross progeny, we individually cultured 100 F2 animals and observed F3 and F4 progeny to specifically identify recombinant populations that failed to produce dead embryos or those that starved at the same rate as VC20019 controls. Positively identified populations were combined for MIP-MAP analysis (**Methods**). The primary candidate mutation for VC40788 is a G405R mutation in the mitochondrial protein B0303.3, which is predicted to have multiple functions including an acetyl-CoA C-acyltransferase activity. *B0303*.*3* has no reported hypomorphic or null mutant alleles but is reported to have an embryonic lethal phenotype by RNAi (Gönczy et al., 2000; Sönnichsen et al., 2005) and its human ortholog *HADHB* is implicated in trifunctional protein deficiency phenotype (Purevsuren et al., 2009; Spiekerkoetter et al., 2003). The identification of a maternal hypomorphic allele of *B0303*.*3* provides a means with which to study this disease and its phenotypes in a nematode model.

**Table 3.** Mapping data summary.

| Strain | Pools | Mean FCR | Class | Mapping Interval | Coding alleles | Likely Candidate |
| --- | --- | --- | --- | --- | --- | --- |
| VC20019 | All but M1 | 0.136 | 3 | -- | -- | -- |
| VC30079 | M5, M6 | -0.740 | 1 | II:7.49-11.5 Mb | 3 | *hpo-35* |
|  |  |  |  | III:5.8-7.6 Mb | 3 | *dig-1* |
| VC30188 | M5, M6 | -1.038 | 1 | II:6.2-12.1 Mb | 1 | *mel-11* |
| VC40196 | M1, M3 | -- | 0 | IV:8.4-13.9 Mb | 13 | -- |
| VC40296 | M5, M6 | -- | 0 | IV:4.2-6.4 Mb | 2 | *rme-2* |
| VC40545 | M1, M3 | -- | 0 | II:4.4-8.1 Mb | 12 | *tsn-1* |
| VC40611 | M1, M3 | -- | 0 | II:6.3-8.1 Mb | 7 | -- |
| VC40788 | M1, M3 | -- | 0 | III:7.6-10.8 Mb | 2 | *B0303.3* |

## Discussion

With advances in sequencing, genome-editing, and imaging, one remaining bottleneck in the characterization of the *C. elegans* genome is our ability to identify the phenotypes associated with gene function (Granier and Vile, 2014; Houle et al., 2010). The ability to quantify population fitness along a spectrum provides a window into gene functions that may otherwise be overlooked under current experimental paradigms. Dissecting the contribution of weaker alleles will help to generate new gene networks and build upon our understanding of worm development, reproduction, and overall fitness. With PhenoMIP, we analysed strains from the Million Mutation Project, which offers a unique library of mutagenized genomes with coding and non-coding elements that remain largely unexplored. We efficiently identified phenotypic traits related to population fitness in a high-throughput manner by pooling multiple MMP strains in a multi-generational experiment and sequencing these populations with molecular inversion probes.

To use MIPs as a means of barcoding strains for population analysis, we designed a series of probes for the 2007 MMP strains and tested a subset on the MMP collection. We observed that we could accurately gauge a strain’s relative abundance within a sample. By sequencing multiple genomic mixtures, we confirmed a low false positive rate, suggesting we could use MIPs to accurately identify subpopulations with abundance as low as 8×10^−4^ which translates to better than 1 in 1000 genomes per sample.

As a demonstration of this method, we pooled MMP strains into groups and dissected population composition over multiple generations. Our observations suggest that this form of population barcoding is indeed capable of identifying specific Million Mutation Project strains with differing levels of relative fitness. Our analysis shows that PhenoMIP identifies reproducible condition-dependent population stratification among populations that have been separated for multiple generations. Based on the strains tested thus far, we estimate upwards of 82% of MMP strains may harbour alleles contributing to fitness phenotypes in the range of class 0 to class 2. Given the mutagenized and inbred nature of the MMP strains (Thompson et al., 2013), it is not surprising to find such an array of fitness phenotypes. These strains, however, represent a valuable resource to study fitness as the causative alleles of these effects may be in putative essential genes, poorly characterized genes with only small effects on fitness, or even regulatory regions of the genome.

The observed population-level phenotypes presented in this work are a readout of relative fitness in a multi-strain competitive environment. Depending on selection and pooling method, weaker changes to relative fitness may be attributed to the population mixture rather than the selection variable itself. For instance, in our series of experiments, pools were initially generated by combining small numbers of larval animals as a seeding parental population that was expanded before aliquoting out to replicate experiments. During the initial expansion of the seed population, the stochastic loss of even a single parental animal could impact the abundance of a strain in the initial stages of the experiment. Conversely, we saw in our analysis of pool M8, that the doubling of VC20019 animals in the initial pooling also affected the population structure and mean fold-change rate of VC20019 itself. A potential solution to mitigate “seeding” variation would be to bleach synchronize (Stiernagle, 2006) all of the target strains to the L1 larval stage and then combine them in equal portions into a single population before aliquoting to replicate experiments. Another influence on population structure is the group of Class 4 strains identified in our study. Their rapid growth and expansion can lead to drastic population stratification and the premature loss of subpopulations. In these cases, the quantitative phenotyping of less fit strains may be hindered, less informative, or potentially less accurate when analysing a multi-generational experiment. Therefore, depending on the nature of the experiment, it may be more advantageous to consider pooling strains of a similar fitness based on prior phenotype data. Our observations also suggest that food source can alter population growth with food scarcity contributing to greater variation between replicates. For example, our OP50 replicates may have experienced premature starvation or uneven food distribution amongst populations, leading to lower population sizes and possibly affecting the consistency of the OP50-grown replicates. For an auxotrophic food source such as OP50, it would be best to highly concentrate cultures in order to generate a thicker lawn for nematode populations to consume. Lastly, the method and timing of population transfer is a potential source of selective influence. Our data suggested that chunking versus washing populations to propagate them did introduce technical variation with some strains. A method of population transfer that was not addressed in this work is the bleach synchronization method (Stiernagle, 2006), which would add the benefit of removing sporadic contamination while indirectly assaying developmental timing and fecundity. Some strains may also be differentially sensitive to bleaching, starvation or recovery from starvation (Baugh, 2013; Webster et al., 2019). Over many generations, the above technical variation can amplify within the population, potentially skewing the changes observed. Therefore, when applying specific selective pressures to a population (temperature, food source, RNAi, etc.), the proper use of control conditions and replicates can help to reduce the effects of technical variation with minimal impact to the sequencing burden of the experiment.

Looking to the future, given the wide range of sequenced strains available from the Million Mutation Project and *Caenorhabditis elegans* Natural Diversity Resource (Cook et al., 2017), a more extensive competitive fitness assessment by PhenoMIP would set the stage for generating balanced pools of strains based on similar growth rates. From similarly profiled strains, balanced pools could be generated randomly or based on parameters such as geographic distribution or specific genotypes or haplotypes of interest. These pools could be used to screen for phenotypic differences among any number of conditions from temperature or food source (Dirksen et al., 2016; Zhang et al., 2017) to resource limitation, small molecule exposure, or pathogen infection. Recently, Webster et al., utilized RAD-seq techniques to assess starvation resistance on a multiplexed pool of 96 wild isolate strains (Webster et al., 2019). This form of competitive fitness selection is an ideal experimental context for PhenoMIP to increase potential throughput by addressing additional parameters or variables related to starvation response. Furthermore, the process of pooled competition facilitates screening on multiple strains in scenarios where the substrates or reagents to test have limited availability. In combination with GWAS and genetic mapping, PhenoMIP could prove useful in assembling a greater understanding of the many unexplored gene and regulatory sequence functions within the *C. elegans* genome.

To our knowledge these experiments are the first to use molecular inversion probes to analyse *C. elegans* populations for relative fitness. With PhenoMIP, we analysed 217 MMP strains across 95 replicate conditions and 29 timepoints for a total of 393 genomic samples. A similar analysis of our experimental data via whole genome sequencing across 393 genomic samples would be prohibitively expensive. In contrast, our data can be generated on the equivalent of a single Illumina NextSeq run. Targeted sequencing by PhenoMIP permits experimentation at a scale well beyond what is reasonably accomplished by standard WGS. PhenoMIP, however, is not without its caveats as the data generated is limited to assessing relative abundance and the variants assessed are limited to the population of strains in the experiment. We believe, however, that the initial processing steps and costs as well as the “limited” variant diversity of the data are outweighed by the increase in experimental throughput.

PhenoMIP has the potential to be applied beyond the MMP and wild isolate strains to the quantitative analyse of genomic variants in many contexts. Coupled with genome-level editing techniques, PhenoMIP could be useful in studying allelic series or mutants of entire pathways for subtle phenotypic effects. The assay format could be converted to look at selection of phenotypes occurring within a single event or generation, as in a bulk taxis assay or as a method for targeted genome monitoring under selective conditions. The fundamental leverage of this method is the use of MIPs to reduce the sequencing burden while maintaining informative parity with WGS formats in identifying subpopulation frequency. In doing so, the throughput of experimentation can be increased without raising experimental sequencing costs.

## Methods

### MIP site selection and design

MIP sites were selected in two rounds. Initially the entire MMP SNV data set was used to select for sites that were spaced a minimum of 300 bp apart to avoid potential collisions with neighbouring probes. Site selection and rejection was completed in a linear manner based on the first available SNV on each linkage group within the data set. Locations were not filtered or optimized to reduce the occurrence of neighbouring SNVs within the 300 bp window. The initial set of MMP mutant strain MIP sites was then used to remove candidate sites from the MMP wild isolate data set. Any wild isolate sites within a 350 bp window of mutant candidate sites was removed from selection. Of the remaining wild isolate SNV sites, a 350 bp selection window was used to identify potential MIP sites. The list of candidate MIP sites were used to design and score MIPs based on previously published criteria (Mok et al., 2017). The list of designed MIPS was subdivided into each individual strain where the highest-scoring MIP for each linkage group was identified. Of the six MIPs designed for each strain, four were randomly selected for use in population analysis (**Supplemental Data SD1**)

### MIP library pooling, preparation and sequencing

MIPs were pooled based on worm pools being tested and generated as previously published (Mok et al, 2017). Individual MIPs were normalized to a concentration of 100 uM and pooled to a maximum volume of 85 ul. 10 ul of 10X Polynucleotide Kinase (PNK) Buffer and 5 ul of PNK were added to a volume of 85 ul pooled MIPs before incubating for 45 minutes at 37°C and 20 minutes at 80°C. This pool was then diluted to a working concentration of 330 nM. MIP libraries were generated with 500 ng genomic DNA and appropriate MIP pools as previously described in Mok et al., 2017. Libraries were sequenced on Illumina MiSeq or NextSeq systems. Libraries across pools ranged between 8.3×10^6^ and 32.7×10^6^ total reads with an average 1507 reads per probe.

### Worm maintenance and pooling

Worms were maintained on standard nematode growth media (NGM) seeded with OP50. Worm pools were generated from well-fed source plates using exclusively twenty L1 or L4 animals for each strain. Starting pools were grown on 15cm NGM made with 8X peptone and seeded with NA22 or HT115. Pools were grown at 20°C to starvation as mostly L1 animals (96-120 hours) before washing off with 10-15 mL M9. Worms were pelleted and aspirated to 5-6 mL before population density was assessed. 50-100 ul of pellet was frozen as a representative sample of the initial pooled population. Pools were then redistributed in equal-sized populations between 5000 and 10000 animals on 15 cm NGM plates that were prepared based on experimental conditions and grown for 4 days before being transferred to replicate condition plates either by chunking or washing again. Any remaining animals were washed from plates with double-distilled water, pelleted, and frozen as samples for later analysis. Each cycle of transfer approximately followed a single generation and pooling experiments were propagated for 6-10 generations. Heavily contaminated plates/conditions were terminated from propagation and removed from analysis.

### Mapping of mutant strains

Mutant strains were mapped using either the VC20019 mapping strain or DM7448 (VC20019; Ex[pmyo-3::YFP]). Briefly, mapping strain males were crossed with mutant hermaphrodites. 15-20 cross progeny L4 hermaphrodites were selected to a single 10 cm OP50-seeded NGM plate and grown to starvation before propagating a subpopulation to a replicate 10 cm plate. Slow growth mutants were mapped on 10 cm NGM plates seeded with OP50 and grown at 20°C. Mapping populations were propagated under selection for four to seven generations. Representative samples were chosen to extract genomic DNA as template for MIP-MAP libraries and then sequenced on Illumina MiSeq or NextSeq instruments. MIP-MAP analysis was completed as previously described (Mok et al., 2017).

### Competitive Fitness MIP library data analysis

For each specific MIP pool, reads were initially analysed as previously described (Mok et al., 2017) with the exclusion of the normalization step for each MIP. After abundance of each MIP was calculated, an average abundance was calculated for each strain as well as a standard deviation across this average. These values were used in downstream analysis of population structure across multiple timepoints.

Population structure and fold-change analysis was calculated across each experiment using the amalgamated data from above. Strains with a starting abundance value below 2.5×10^−3^ were eliminated from downstream population analysis. Remaining data were further transformed with any values below 1.0×10^−3^ being converted to this value to accommodate log growth analysis. Total fold-change and mean fold change are calculated based on starting and end-point changes in abundance versus total generations (one generation per expansion). In samples with negative trajectories, however, the final generation of growth was calculated as the first instance of abundance at or below the lower limit of 1.0×10^−3^. Mean fold-change rate was calculated based on the total fold-change abundance in the final generation of growth divided by the expected number of generations passed.

### Data Availability

File SD1 contains molecular inversion probe sequences and data for all 2007 MMP strains and 40 wild isolates of the Million Mutation Project. Four candidate probes for each strain were designed and listed in this file. File SD2 contains all information used in the false positive and precision analysis of PhenoMIP. File SD3 contains all mean FCR data for each strain on each replicate in each experimental pool. Custom scripts used to analyse sequencing data are available upon request. Raw sequence files for each pool are available upon request.

## Acknowledgements

Some strains were provided by the CGC, which is funded by the NIH Office of Research and Infrastructure Programs (P40 OD010440). Work by C.M was supported by the Canadian Institutes of Health Research MFE-135408. Work in the R.H.W laboratory was supported by an ARRA GO grant HG005921 from the NHGRI, an R21 grant HG007201 from the NIH, and by the William H. Gates Chair of Biomedical Sciences. The work in the D.G.M laboratory was supported by the Canadian Institutes of Health Research.

## Supplemental Figure Legends

**Supplemental Figure S1.**
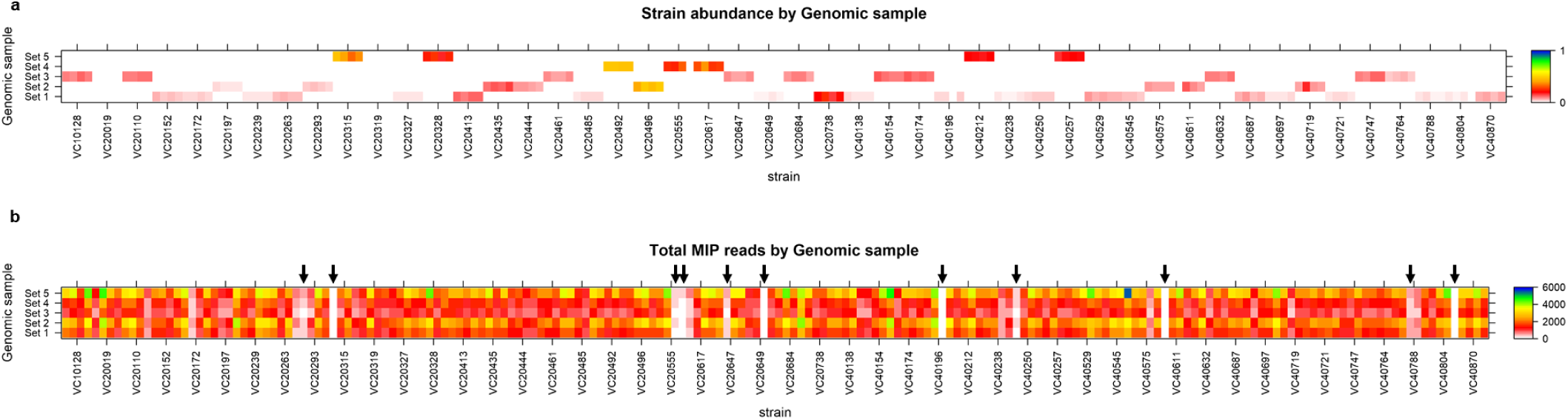
MIPs provide sufficient read depth to specific subpopulations of strain abundance in complex compositions of genomic DNA. (a) Strains from non-overlapping sets of mixed genomic samples are identified using a multiplexed pool of MIPs. Strain abundance for each set is indicated by the heatmap legend. (b) A heatmap of total reads per MIP per set broken down by specific strain with black arrows indicating probes with total reads below 20% of the mean read depth across the set.

**Supplemental Figure S2.**
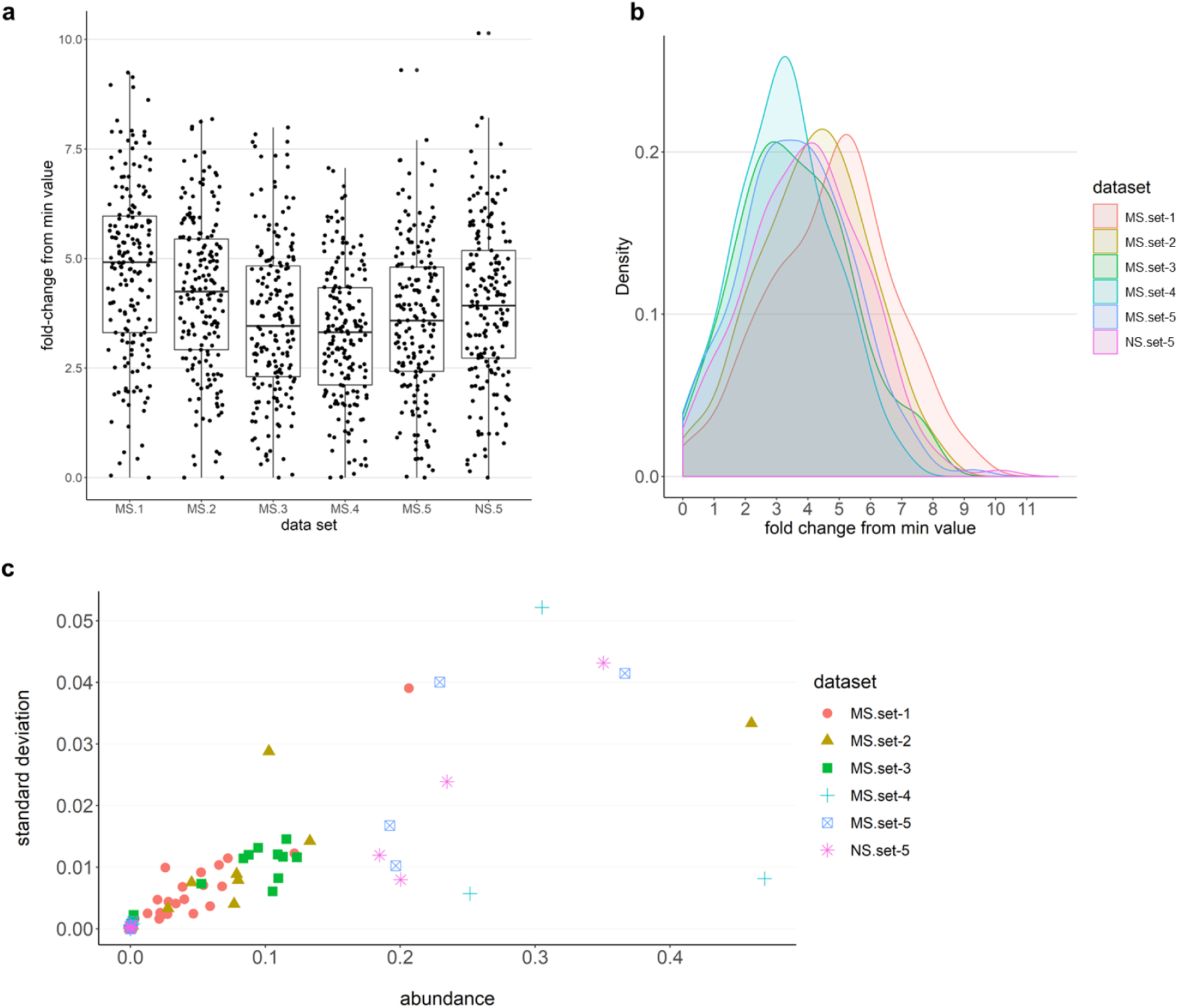
MIP pooling across multiple targets remains balanced and precise. (a) Boxplot of sequencing libraries for the same set of probes across 5 separate genomic templates overlaid with the fold-change for each probe based on the probe with the fewest reads in each set. (b) a kernel density plot of each dataset based on the fold-change in read depth of each probe (MS = MiSeq-generated data; NS = NextSeq-generated data). (c) A scatterplot of abundance for all strains within each sequenced set versus the standard deviation of the 3 to 4 probes used to calculate that abundance.

**Supplemental Figure S3.**
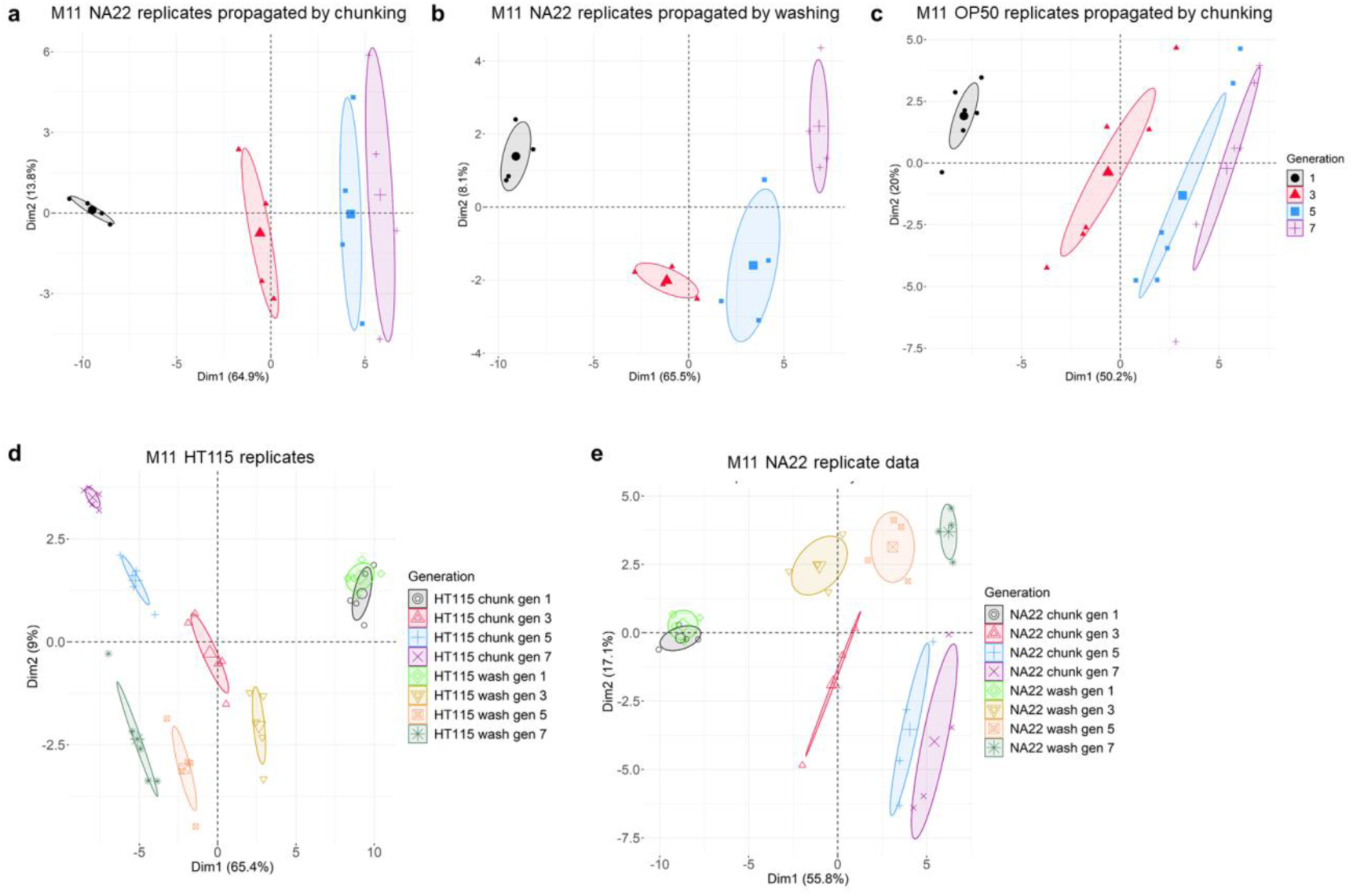
Principal component analysis of M11 samples suggest consistent changes to population structure at each generation. PCA of M11 datasets separated by combined food source and transfer method into (a) NA22 replicates propagated by chunking, (b) NA22 replicates propagated by washing and (c) OP50 replicates propagated by chunking. PCA of M11 HT115 replicate (d) and NA22 replicate (e) data projected along principal components 1 and 2 with samples identified by combination of transfer method (chunking or washing) and sample generation (1, 3, 5, or 7).

**Supplemental Figure S4.**
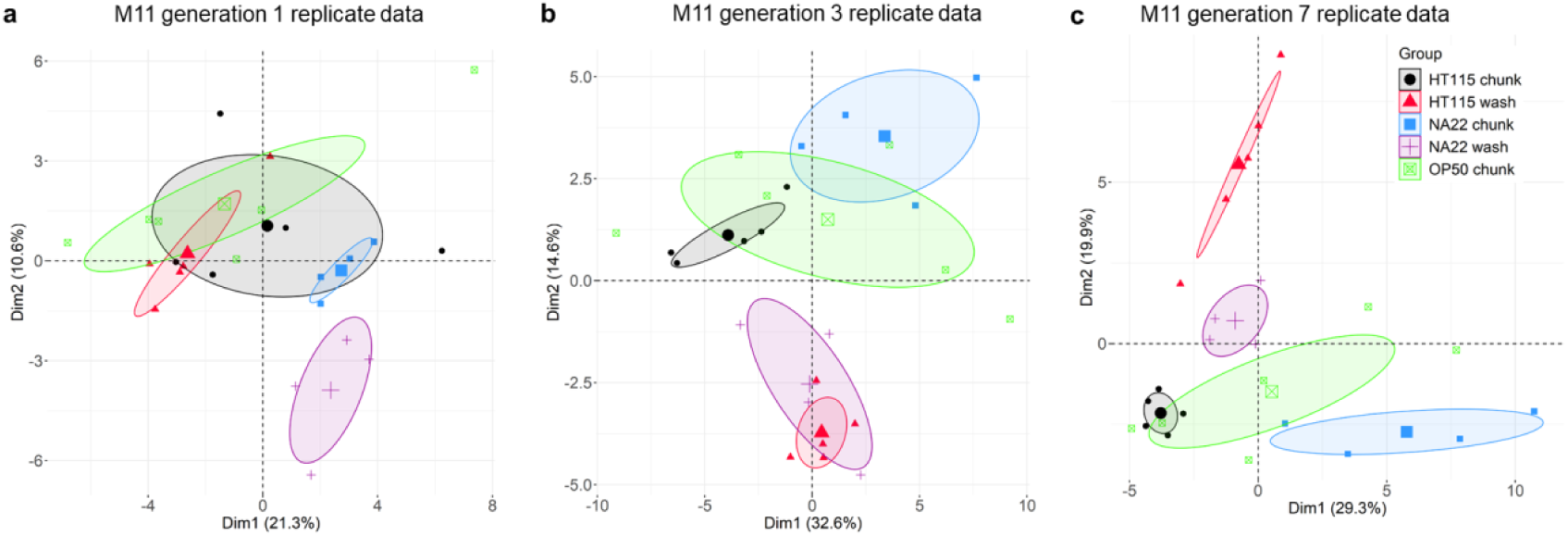
Principal component analysis of M11 samples suggest condition-dependent population structure. PCA of M11 replicate datasets separated into (a) generation 1, (b) generation 3, and (c) generation 7. Samples are projected along principal components 1 and 2 for each individual data set and identified by combination of food source and transfer method.

**Supplemental Figure S5.**
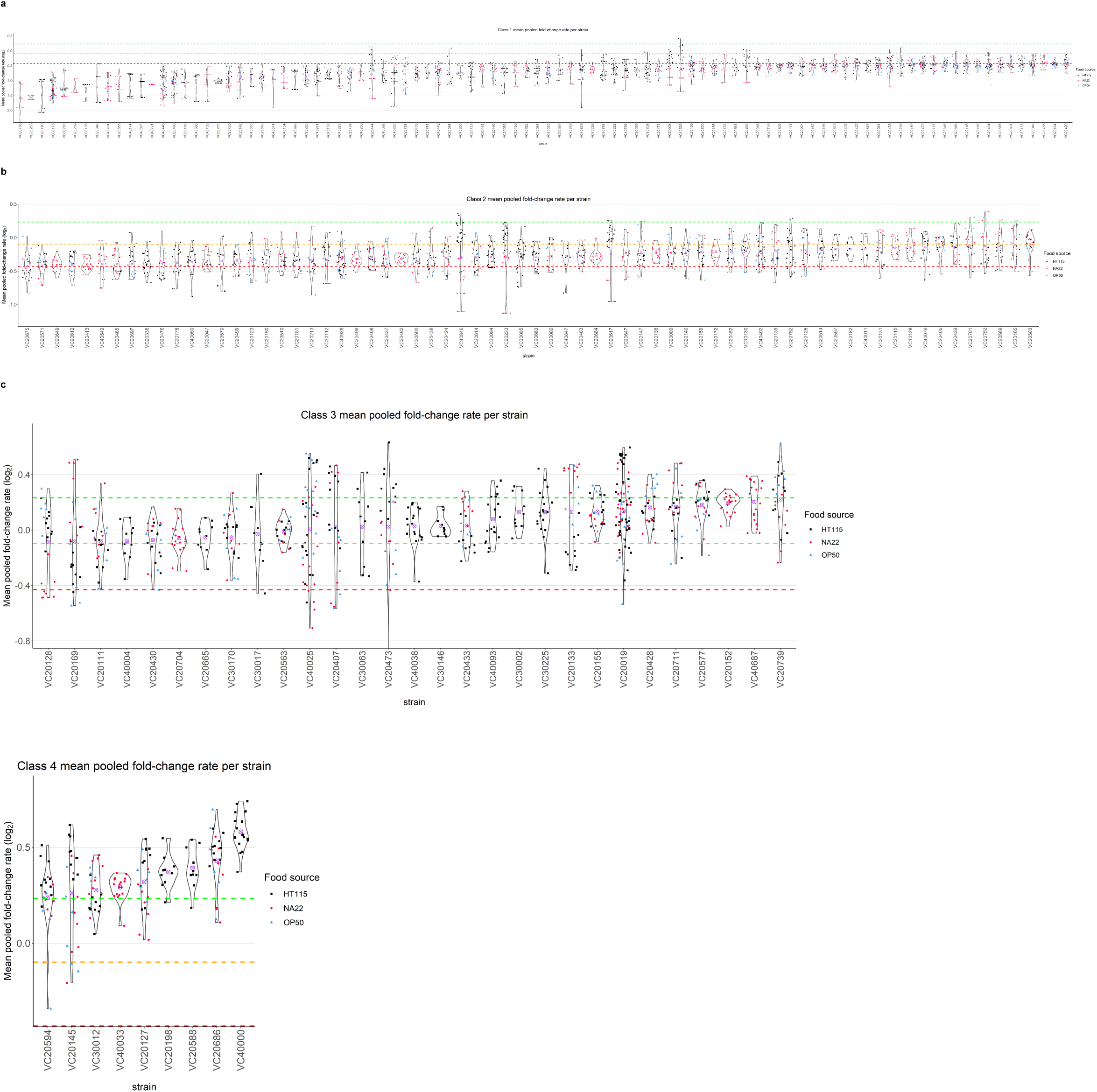
PhenoMIP can generate a gradient of fitness phenotypes from severe to subtle. Violin plots of mean fold change rate per replicate for 202 MMP strains across the 4 defined classes 1 (a), class 2 (b), class 3 (c), and class 4 (d). Each violin plot discriminates between food sources HT115 (black squares), NA22 (red circles) and OP50 (blue triangles). Strains are sorted within class by the mean FCR (log_2_, purple cross) of all replicate conditions for that strain. Coloured dotted lines represent category boundaries using an FCR of -0.4315 (red), -0.0.985 (yellow), and 0.2327 (green).

**Supplemental figure S6.**
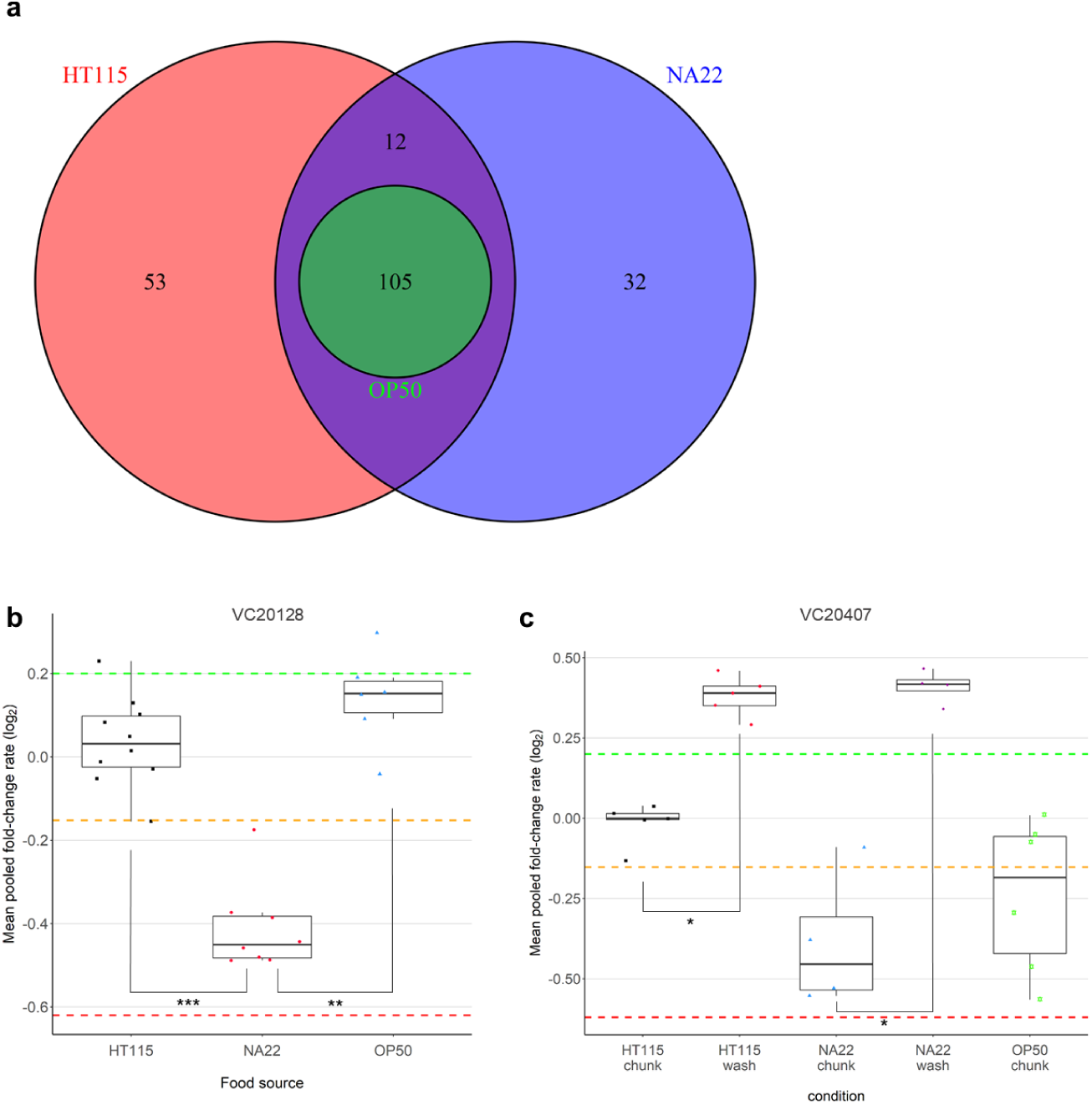
MMP strains were tested across a trio of food sources. (a) Venn diagram of each food source used in the PhenoMIP assays and the number of strains tested with HT115 (pink), NA22 (purple) and OP50 (green). 105 strains were tested on all three food conditions. (b) VC20128 data from the same pool (M10) suggests specific fitness differences between growth on NA22 versus growth on HT115 and OP50. (c) VC20407 data from the same pool (M11) suggests significant changes to growth when comparing samples transferred by chunking versus washing – regardless of food source. Coloured dotted lines represent category boundaries using an FCR of -0.4315 (red), -0.0.985 (yellow), and 0.2327 (green). * p < 0.05; ** p < 0.01; *** p < 0.001 by Kruskal-Wallis with p-values adjusted for multiple testing by Benjamini-Hochberg method

**Supplemental Figure S8.**
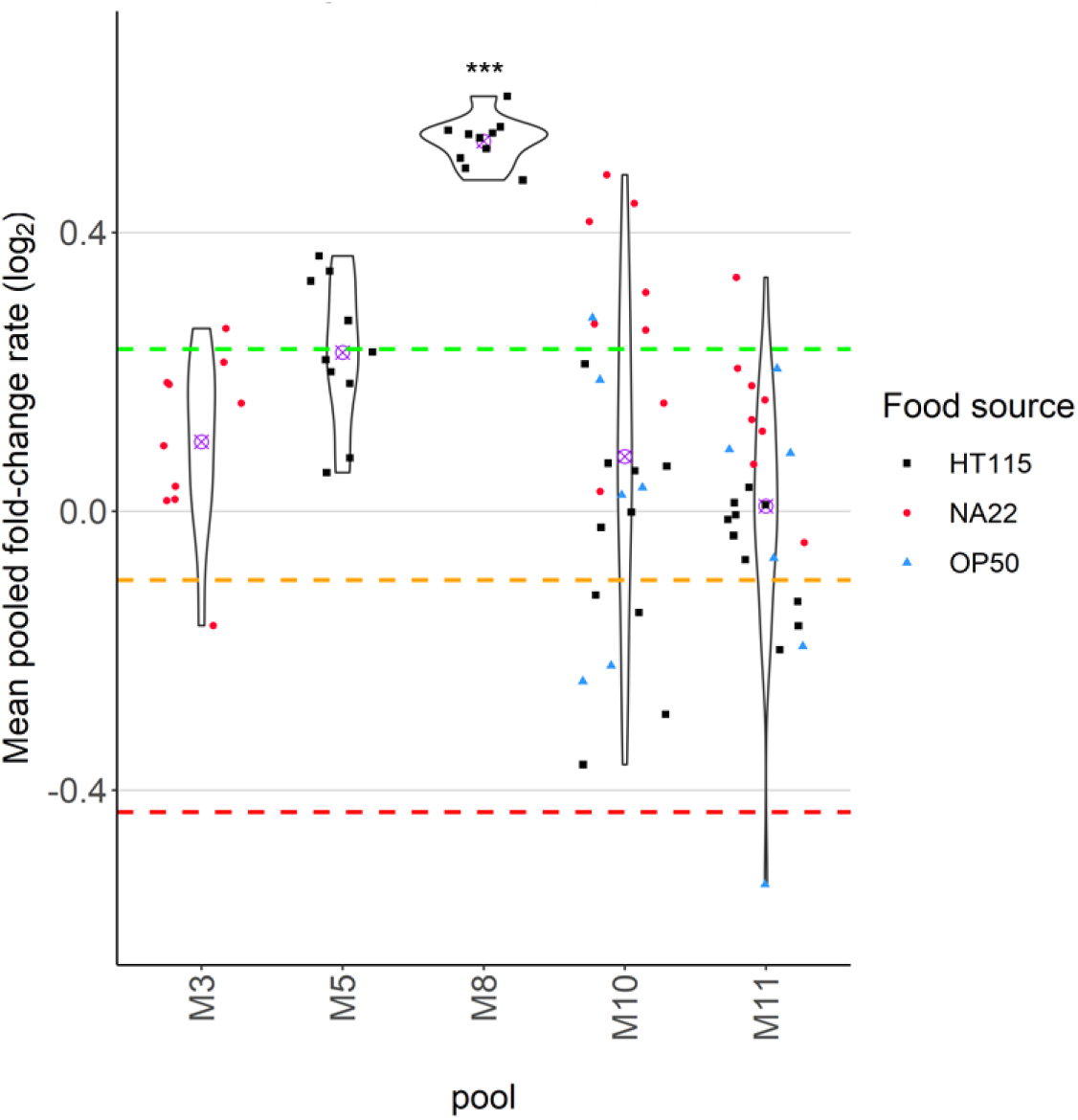
Violin plots of VC20019 mean FCR for all replicates grouped by pool. Violin plots for VC20019 replicates in each pool were generated with M11 represented by combining datasets based on food source (HT115 = HT115 chunk + HT115 wash; NA22 = NA22 chunk + NA22 wash). M8 replicate data is significantly different compared to M3, M5, M10 and M11 which are not significantly different from each other. Each violin plot discriminates between food sources HT115 (black squares), Na22 (red circles) and OP50 (blue triangles) and mean FCR (purple cross). Coloured dotted lines represent category boundaries using an FCR of -0.4315 (red), -0.0.985 (yellow), and 0.2327 (green). *** p < 0.001 by Kruskal-Wallis with p-values adjusted for multiple testing by Benjamini-Hochberg method.

**Supplemental Figure S9.**
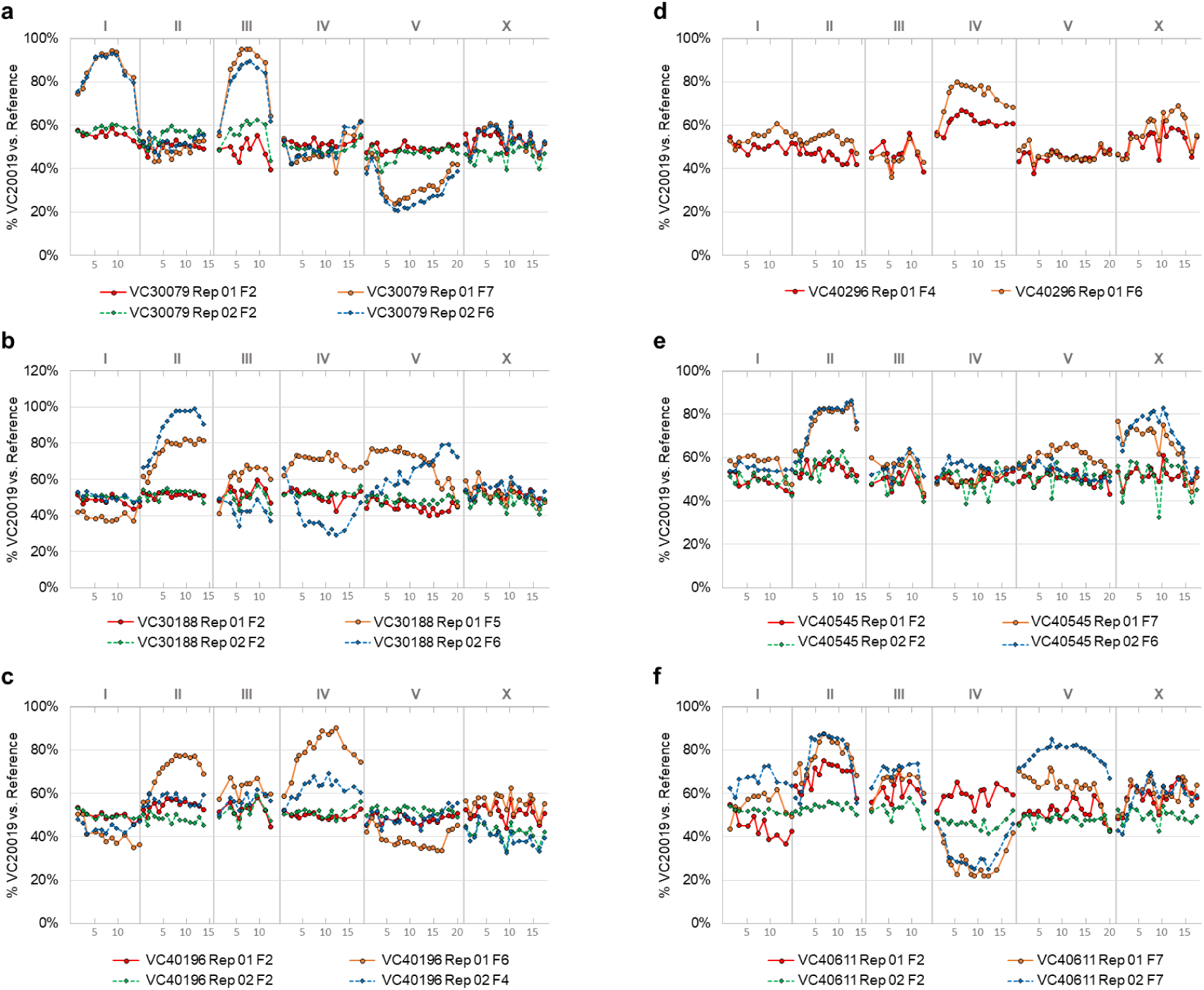
MIP-MAP data for 6 strains categorized as class 0 or class 1 by mean FCR. Strains were mapped using VC20019 with the y-axis representing the proportion of VC20019 present versus all reads for a MIP target at each locus across the genome. Strains were mapped in replicate (solid versus dotted lines) and sequenced at two timepoints each (ie. F2 vs F4). The strains mapped in this fashion were (a) VC30079, (b) VC30188, (c) VC40196, (d) VC40296, (e) VC40545, and (f) VC40611. X-axis units are in megabases across each chromosome.

**Supplemental Table S1.** Class 0 mutants, not analysed due to low abundance at experimental start.

| Strain | Pool(s) | Initial abundance |
| --- | --- | --- |
| VC20190 | M5 | 0.00180 / 0.00077 |
| VC20245 | M1 / M3 | 0.00027 / 0.00026 |
| VC20262 | M5 | 0.00074 / 0.00128 |
| VC20315 | M1 / M3 | 0.00000 / 0.00059 |
| VC20328 | M1 / M3 | 0.00136 / 0.00216 |
| VC20338 | M5 | 0.00166 / 0.00209 |
| VC40291 | M1 / M3 | 0.00114 / 0.00000 |
| VC40296 | M5 | 0.00118 / 0.00017 |
| VC40545 | M1 / M3 | 0.00064 / 0.00017 |
| VC40611 | M1 / M3 | 0.00000 / 0.00034 |
| VC40697 | M1 / M3 | 0.00176 / 0.00219 |
| VC40745 | M5 | 0.00098 / 0.00118 |
| VC40747 | M1 / M3 | 0.00151 / 0.00147 |
| VC40788 | M1 / M3 | 0.00038 / 0.00029 |
| VC40804 | M1 / M3 | 0.00054 / 0.00036 |

## Literature Cited

Araya CL, Kawli T, Kundaje A, Jiang L, Wu B, Vafeados D, Terrell R, Weissdepp P, Gevirtzman L, MacE D, Niu W, Boyle AP, Xie D, Ma L, Murray JI, Reinke V, Waterston RH, Snyder M. 2014. Regulatory analysis of the C. Elegans genome with spatiotemporal resolution. Nature. doi:10.1038/nature13497

Baugh LR. 2013. To grow or not to grow: Nutritional control of development during Caenorhabditis elegans L1 Arrest. Genetics. doi:10.1534/genetics.113.150847

Boeck ME, Huynh C, Gevirtzman L, Thompson OA, Wang G, Kasper DM, Reinke V, Hillier LW, Waterston RH. 2016. The time-resolved transcriptome of C. Elegans. Genome Res 26:1441–1450. doi:10.1101/gr.202663.115

Brenner S. 1974. The Genetics of Caenorhabditis elegans. Genet. doi:10.1111/j.1749-6632.1999.tb07894.x

Cao J, Packer JS, Ramani V, Cusanovich DA, Huynh C, Daza R, Qiu X, Lee C, Furlan SN, Steemers FJ, Adey A, Waterston RH, Trapnell C, Shendure J. 2017. Comprehensive single-cell transcriptional profiling of a multicellular organism. Science (80-). doi:10.1126/science.aam8940

Cook DE, Zdraljevic S, Roberts JP, Andersen EC. 2017. CeNDR, the Caenorhabditis elegans natural diversity resource. Nucleic Acids Res. doi:10.1093/nar/gkw893

Crombie TA, Saber S, Saxena AS, Egan R, Baer CF. 2018. Head-to-head comparison of three experimental methods of quantifying competitive fitness in C. elegans. PLoS One 13:1–11. doi:10.1371/journal.pone.0201507

Daugherty AC, Yeo RW, Buenrostro JD, Greenleaf WJ, Kundaje A, Brunet A. 2017. Chromatin accessibility dynamics reveal novel functional enhancers in C. elegans. Genome Res. doi:10.1101/gr.226233.117

De-Souza EA, Camara H, Salgueiro WG, Moro RP, Knittel TL, Tonon G, Pinto S, Pinca APF, Antebi A, Pasquinelli AE, Massirer KB, Mori MA. 2019. RNA interference may result in unexpected phenotypes in Caenorhabditis elegans. Nucleic Acids Res. doi:10.1093/nar/gkz154

De Stasio EA, Dorman S. 2001. Optimization of ENU mutagenesis of Caenorhabditis elegans. Mutat Res - Genet Toxicol Environ Mutagen. doi:10.1016/S1383-5718(01)00198-X

Diaz SA, Viney M. 2014. Genotypic-specific variance in Caenorhabditis elegans lifetime fecundity. Ecol Evol. doi:10.1002/ece3.1057

Dirksen P, Marsh SA, Braker I, Heitland N, Wagner S, Nakad R, Mader S, Petersen C, Kowallik V, Rosenstiel P, Félix MA, Schulenburg H. 2016. The native microbiome of the nematode Caenorhabditis elegans: Gateway to a new host-microbiome model. BMC Biol 14:1–16. doi:10.1186/s12915-016-0258-1

Doitsidou M, Poole RJ, Sarin S, Bigelow H, Hobert O. 2010. C. elegans mutant identification with a one-step whole-genome-sequencing and SNP mapping strategy. PLoS One 5. doi:10.1371/journal.pone.0015435

Elvin M, Snoek LB, Frejno M, Klemstein U, Kammenga JE, Poulin GB. 2011. A fitness assay for comparing RNAi effects across multiple C. elegans genotypes. BMC Genomics. doi:10.1186/1471-2164-12-510

Fraser a G, Kamath RS, Zipperlen P, Martinez-Campos M, Sohrmann M, Ahringer J. 2000. Functional genomic analysis of C. elegans chromosome I by systematic RNA interference. Nature 408:325–330. doi:10.1038/35042517

Fraser AG. 2000. Functional genomic analysis of C. elegans chromosome I by systematic RNA interference. Nature 408:325–330.

Gönczy P, Echeverri C, Oegema K, Coulson A, Jones SJM, Copley RR, Duperon J, Oegema J, Brehm M, Cassin E, Hannak E, Kirkham M, Pichler S, Flohrs K, Goessen A, Leidel S, Alleaume AM, Martin C, Özlü N, Bork P, Hyman AA. 2000. Functional genomic analysis of cell division in C. elegans using RNAi of genes on chromosome III. Nature. doi:10.1038/35042526

Gracida X, Calarco JA. 2017. Cell type-specific transcriptome profiling in C. elegans using the Translating Ribosome Affinity Purification technique. Methods. doi:10.1016/j.ymeth.2017.06.023

Granier C, Vile D. 2014. Phenotyping and beyond: Modelling the relationships between traits. Curr Opin Plant Biol. doi:10.1016/j.pbi.2014.02.009

Hardenbol P, Banér J, Jain M, Nilsson M, Namsaraev EA, Karlin-Neumann GA, Fakhrai-Rad H, Ronaghi M, Willis TD, Landegren U, Davis RW. 2003. Multiplexed genotyping with sequence-tagged molecular inversion probes. Nat Biotechnol. doi:10.1038/nbt821

Hiatt JB, Pritchard CC, Salipante SJ, O’Roak BJ, Shendure J. 2013. Single molecule molecular inversion probes for targeted, high-accuracy detection of low-frequency variation. Genome Res 23:843–854. doi:10.1101/gr.147686.112

Houle D, Govindaraju DR, Omholt S. 2010. Phenomics: The next challenge. Nat Rev Genet. doi:10.1038/nrg2897

Jaramillo-Lambert A, Fuchsman AS, Fabritius AS, Smith HE, Golden A. 2015. Rapid and Efficient Identification of Caenorhabditis elegans Legacy Mutations Using Hawaiian SNP-Based Mapping and Whole Genome Sequencing. G3 5:1007–1019. doi:10.1534/g3.115.017038

Jorgensen EM, Mango SE. 2002. The art and design of genetic screens: Caenorhabditis elegans. Nat Rev Genet. doi:10.1038/nrg794

Kaletsky R, Yao V, Williams A, Runnels AM, Tadych A, Zhou S, Troyanskaya OG, Murphy CT. 2018. Transcriptome analysis of adult Caenorhabditis elegans cells reveals tissue-specific gene and isoform expression. PLoS Genet. doi:10.1371/journal.pgen.1007559

Kamath RS, Fraser AG, Dong Y, Poulin G, Durbin R, Gotta M, Kanapin A, Le Bot N, Moreno S, Sohrmann M, Welchman DP, Zipperlen P, Ahringer J. 2003. Systematic functional analysis of the Caenorhabditis elegans genome using RNAi. Nature 421:231–7. doi:10.1038/nature01278

Kevin S, Barstead RJ, Moerman DG. 2006. *C. elegans* Deletion Mutant ScreeningC. Elegans. doi:10.1385/1-59745-151-7:51

Lehner B, Crombie C, Tischler J, Fortunato A, Fraser AG. 2006. Systematic mapping of genetic interactions in Caenorhabditis elegans identifies common modifiers of diverse signaling pathways. Nat Genet 38:896–903. doi:10.1038/ng1844

Minevich G, Park DS, Blankenberg D, Poole RJ, Hobert O. 2012. CloudMap: A cloud-based pipeline for analysis of mutant genome sequences. Genetics 192:1249–1269. doi:10.1534/genetics.112.144204

Mok CA, Au V, Thompson OA, Edgley ML, Gevirtzman L, Yochem J, Lowry J, Memar N, Wallenfang MR, Rasoloson D, Bowerman B, Schnabel R, Seydoux G, Moerman DG, Waterston RH. 2017. MIP-MAP: High-throughput mapping of Caenorhabditis elegans temperature-sensitive mutants via molecular inversion probes. Genetics 207:447–463. doi:10.1534/genetics.117.300179

Parrish S, Fleenor J, Xu SQ, Mello C, Fire A. 2000. Functional anatomy of a dsRNA trigger: Differential requirement for the two trigger strands in RNA interference. Mol Cell. doi:10.1016/S1097-2765(00)00106-4

Perez MF, Francesconi M, Hidalgo-Carcedo C, Lehner B. 2017. Maternal age generates phenotypic variation in Caenorhabditis elegans. Nature. doi:10.1038/nature25012

Purevsuren J, Fukao T, Hasegawa Y, Kobayashi H, Li H, Mushimoto Y, Fukuda S, Yamaguchi S. 2009. Clinical and molecular aspects of Japanese patients with mitochondrial trifunctional protein deficiency. Mol Genet Metab. doi:10.1016/j.ymgme.2009.07.011

Ramani AK, Chuluunbaatar T, Verster AJ, Na H, Vu V, Pelte N, Wannissorn N, Jiao A, Fraser AG. 2012. The majority of animal genes are required for wild-type fitness. Cell 148:792–802. doi:10.1016/j.cell.2012.01.019

Richards JL, Zacharias AL, Walton T, Burdick JT, Murray JI. 2013. A quantitative model of normal Caenorhabditis elegans embryogenesis and its disruption after stress. Dev Biol. doi:10.1016/j.ydbio.2012.11.034

Schnabel R, Hutter H, Moerman D, Schnabel H. 1997. Assessing normal embryogenesis in Caenorhabditis elegans using a 4D microscope: Variability of development and regional specification. Dev Biol. doi:10.1006/dbio.1997.8509

Sönnichsen B, Koski LB, Walsh a, Marschall P, Neumann B, Brehm M, Alleaume a-M, Artelt J, Bettencourt P, Cassin E, Hewitson M, Holz C, Khan M, Lazik S, Martin C, Nitzsche B, Ruer M, Stamford J, Winzi M, Heinkel R, Röder M, Finell J, Häntsch H, Jones SJM, Jones M, Piano F, Gunsalus KC, Oegema K, Gönczy P, Coulson a, Hyman a a, Echeverri CJ. 2005. Full-genome RNAi profiling of early embryogenesis in Caenorhabditis elegans. Nature 434:462–9. doi:10.1038/nature03353

Spiekerkoetter U, Sun B, Khuchua Z, Bennett MJ, Strauss AW. 2003. Molecular and phenotypic heterogeneity in mitochondrial trifunctional protein deficiency due to β-subunit mutations. Hum Mutat. doi:10.1002/humu.10211

Stiernagle T. 2006. Maintenance of C. elegans. WormBook.

Sulston JE, Schierenberg E, White JG, Thomson JN. 1983. The embryonic cell lineage of the nematode Caenorhabditis elegans. Dev Biol. doi:10.1016/0012-1606(83)90201-4

Thompson O, Edgley M, Strasbourger P, Flibotte S, Ewing B, Adair R, Au V, Chaudhry I, Fernando L, Hutter H, Kieffer A, Lau J, Lee N, Miller A, Raymant G, Shen B, Shendure J, Taylor J, Turner EH, Hillier LW, Moerman DG, Waterston RH. 2013. The million mutation project: A new approach to genetics in Caenorhabditis elegans. Genome Res 23:1749–1762. doi:10.1101/gr.157651.113

Warf MB, Shepherd BA, Johnson WE, Bass BL. 2012. Effects of ADARs on small RNA processing pathways in C. elegans. Genome Res. doi:10.1101/gr.134841.111

Warner AD, Gevirtzman L, Hillier LDW, Ewing B, Waterston RH. 2019. The C. elegans embryonic transcriptome with tissue, time, and alternative splicing resolution. Genome Res. doi:10.1101/gr.243394.118

Webster AK, Hung A, Moore BT, Guzman R, Jordan JM, Kaplan REW, Hibshman JD, Tanny RE, Cook DE, Andersen E, Baugh LR. 2019. Population Selection and Sequencing of Caenorhabditis elegans Wild Isolates Identifies a Region on Chromosome III Affecting Starvation Resistance. G3&amp;#58; Genes|Genomes|Genetics. doi:10.1534/g3.119.400617

Wormbase web site. 2019. release WS271, 15 March 2019. Wormbase. http://www.wormbase.org

Zhang F, Berg M, Dierking K, Félix MA, Shapira M, Samuel BS, Schulenburg H. 2017. Caenorhabditis elegans as a model for microbiome research. Front Microbiol. doi:10.3389/fmicb.2017.00485

